# Gene Expression Risk Scores for COVID-19 Illness Severity

**DOI:** 10.1101/2021.08.24.457521

**Authors:** Derick R Peterson, Andrea M Baran, Soumyaroop Bhattacharya, Angela R Branche, Daniel P Croft, Anthony M Corbett, Edward E Walsh, Ann R Falsey, Thomas J Mariani

**Affiliations:** Department of Biostatistics and Computational Biology, Rochester General Hospital, Rochester, NY, USA; Division of Neonatology and Pediatric Molecular and Personalized Medicine Program, Department of Pediatrics, Rochester General Hospital, Rochester, NY, USA; Division of Infectious Diseases, Department of Medicine, Rochester General Hospital, Rochester, NY, USA; Division of Pulmonary and Critical Care, Department of Medicine, University of Rochester, Rochester General Hospital, Rochester, NY, USA; Department of Medicine, Rochester General Hospital, Rochester, NY, USA

**Keywords:** COVID-19, Severity, RNA sequencing, ICU, Prediction, Molecular Markers

## Abstract

**Background:** The correlates of COVID-19 illness severity following infection with SARS-Coronavirus 2 (SARS-CoV-2) are incompletely understood.

**Methods:** We assessed peripheral blood gene expression in 53 adults with confirmed SARS-CoV-2-infection clinically adjudicated as having mild, moderate or severe disease. Supervised principal components analysis was used to build a weighted gene expression risk score (WGERS) to discriminate between severe and non-severe COVID.

**Results:** Gene expression patterns in participants with mild and moderate illness were similar, but significantly different from severe illness. When comparing severe versus non-severe illness, we identified >4000 genes differentially expressed (FDR<0.05). Biological pathways increased in severe COVID-19 were associated with platelet activation and coagulation, and those significantly decreased with T cell signaling and differentiation. A WGERS based on 18 genes distinguished severe illness in our training cohort (cross-validated ROC-AUC=0.98), and need for intensive care in an independent cohort (ROC-AUC=0.85). Dichotomizing the WGERS yielded 100% sensitivity and 85% specificity for classifying severe illness in our training cohort, and 84% sensitivity and 74% specificity for defining the need for intensive care in the validation cohort.

**Conclusion:** These data suggest that gene expression classifiers may provide clinical utility as predictors of COVID-19 illness severity.

## Introduction

In December 2019 a novel coronavirus, SARS-CoV-2, was identified in China as a cause of severe pneumonia with explosive human-to human transmission [1]. Illness due to SARS-CoV-2 has been designated COVID-19, and on March 11, 2020, the World Health Organization officially declared SARS-CoV-2 a pandemic. To date there have been over 100 million infections and over 2 million deaths globally due to COVID-19. (Source: https://covid19.who.int/) Although most patients experience mild to moderate disease, 5-10% progress to severe or critical illness with severe pneumonia or respiratory failure [2, 3]. Early in the pandemic it became clear that certain underlying chronic medical conditions, and principally age, were key risk factors for severe disease [4, 5]. While severe disease can occur early in illness, a distinct progression to severe illness occurs in some individuals 7-12 days after symptom onset suggesting transition from a viral phase to an inflammatory phase [6]. In addition, some young individuals without co-morbidities have also developed severe illness, highlighting the incomplete understanding of disease pathogenesis due to SARS-CoV-2 infection [7].

Gene expression provides an unbiased measure of the host response to a pathogen on a cellular level. We and others have previously demonstrated the potential for peripheral blood gene expression patterns to classify the ontogeny and severity of viral respiratory illness [8, 9]. We hypothesized that analysis of gene expression in the blood of patients with SARS-CoV2-related COVID-19 might help identify those at greatest risk for severe symptoms and in need of intensive care. Gene expression analysis might also identify pathways underlying disease pathogenesis and suggest new targets amenable to potential therapeutic interventions.

## Methods

### Acute Illness Evaluation

Adults ≥18 years of age, either hospitalized or recruited from the community with symptoms compatible with COVID-19 and documented to have SAR-CoV-2 by PCR, were eligible for the study. Participants with immunosuppression or symptoms onset greater than 28 days prior to admission were excluded. Hospitalized participants were assessed within 24 hours of admission and outpatients were brought to the clinic within 1-2 days of being identified as SARS-CoV-2 positive. Demographic, clinical, radiographic and laboratory information, date of symptom onset and signs and symptoms of the illness were collected. Medication use was recorded with attention to drugs (steroids, leukotriene agonists, hydroxychloroquine, Kaletra, Remdesivir or biologics) that may affect transcriptional profiling.

### Clinical severity assessment

Severity for COVID-19 participants at enrollment and throughout the illness was assessed using a combination of clinical variables (symptoms, physical findings, radiographic and laboratory values) as well as the National Early Warning Score (NEWS) of 7 graded physiological measurements (respiratory rate; oxygen saturation; oxygen supplementation; temperature; blood pressure; heart rate; level of consciousness) [10]. Severe illness was defined as requiring any of the following: ICU care, high flow oxygen, ventilator support, presser support or evidence of new end organ failure. Non-severe illness was defined as illnesses not meeting severe criteria. In addition, a panel of 4 physicians (3 infectious disease and 1 pulmonary critical care) adjudicated individually, then in a live panel discussion, all non-severe illnesses and categorized them as mild or moderate using the NEWS as well as symptoms and physiologic parameters in the context of underlying diseases and baseline oxygen requirements. Participants were followed for the duration of hospitalization and illness, and outcomes were recorded as the highest level of care required or death.

### Sample Collection and Processing

Approximately 3 ml of whole blood was collected in a Tempus™ Blood RNA Tube at the time of enrollment and stored at -80C until the time of processing. Following centrifugation, RNA was isolated from the pellet using the Tempus Spin RNA Isolation Kit using the manufacturer recommended protocol. Total RNA was processed for globin reduction using GLOBINclear Human Kit as described previously [9].

### RNA Sequencing

Globin-reduced RNA was used for transcriptomic profiling by RNA-seq. cDNA libraries were generated using 200 ng of globin-reduced total RNA. Library construction was performed using the TruSeq Stranded mRNA library kit (Illumina, San Diego, CA). cDNA quantity was determined with the Qubit Flourometer (Life Technologies, Grand Island, NY) and quality was assessed using the Agilent Bioanalyzer 2100 (Agilent, Santa Clara, CA). Libraries were sequenced on the Illumina NovaSeq6000 at a target read depth of ∼20 million 1□×□100-bp single end reads per sample. Sequences were aligned against the human genome version hg38 using the Splice Transcript Alignment to a Reference (STAR) algorithm [11], and counts were generated using HTSeq [12]. Raw counts were divided by participant-specific library size (in millions) to yield counts per million (CPM)-normalized expression, borrowing no information across participants, and gene and sample level filtering was performed to remove outlier samples and low expressing genes. Normalized and filtered analytical data sets were log_2_-transformed (after adding a pseudo-count of 1 CPM) prior to analysis. We excluded data from 19,861 genes with uniformly zero reads, leaving a data set comprised of 39,225 genes from 53 participants. Finally, we retained genes that had normalized counts exceeding 1 CPM in greater than 14 participants (the smallest class size). This resulted in an analytical dataset of 14,228 CPM-normalized genes.

### Statistical Methods

Continuous clinical variables were compared by COVID severity levels using the nonparametric Kruskal-Wallis test, and binary variables by Fisher’s exact test. We tested for differential expression by COVID-19 severity using the nonparametric Wilcoxon rank sum test. To allow adjustment for important clinical covariates, we fit semi-parametric Cox proportional hazards models for normalized gene expression as a function of severe vs non-severe COVID-19, adjusted for race, sex, BMI, days since symptom onset, and library size. The Benjamini-Hochberg procedure was used to control the False Discovery Rate (FDR). Pathway analysis of significantly differentially expressed genes was performed using ENRICHR [13].

### Classifier Development and Testing

We used a version of supervised principal components analysis to build a weighted gene expression risk score (WGERS) to discriminate between severe and non-severe COVID-19. Genes were selected based on their univariate AUC and fold-change, as estimated by Hodges-Lehmann median of all pairwise shifts in log expression, with thresholds selected within the inner loop of nested 20-fold cross-validation. The cross-validation sampling strategy was designed to efficiently approximate leaving out 1 subject from each of the 3 sub-levels of severity. The selected genes were standardized to Z-scores with mean 0 and unit variance, and their first PC score was used as the sole predictor for logistic regression. The outer loop of nested cross-validation was used to estimate the ROC and AUC of the adaptive procedure. The nested pooled AUC corresponds with the ROC curve, and compares samples across and within models. The nested stratified AUC only compares samples from the same model, and thus generally is preferable, but it does not correspond with any single ROC curve. A subsequent run of non-nested cross-validation produced the thresholds used to define the gene set for the final risk score. The WGERS is calculated as the linear combination of the standardized, log_2_-transformed genes that meet the chosen thresholds, with coefficients based on the first principal component loading of the genes, scaled by the coefficient from the univariate logistic regression model.

To perform an independent validation of our risk score, we use a dataset from Overmyer et al. [14]. There were some notable differences between the validation data and our training dataset, including a different definition of severity in the outcome (ICU vs non-ICU) and use of a different normalization for the gene expression data (TPM). Of the 18 genes used in our risk score, 2 were missing in the validation data. We imputed data for these 2 genes via multiple linear regression with coefficients estimated by regressing each on the 16 non-missing CPM-normalized log gene expression values in the training data. We standardized the TPM-normalized validation gene expression data using means and SDs estimated from the training data, and then applied the risk score coefficients from the training data to construct a risk score for each validation subject. Apparent miscalibration required choosing a different WGERS threshold for the validation data due to gene expression measures being generally lower in the validation data compared to the training data. An ROC curve with associated AUC was used to assess the performance of the risk score in the validation data.

## Results

Between April 30th and June 29th 2020, 58 participants with PCR documented COVID-19 illnesses were enrolled from inpatient and outpatient settings. Of these, 3 participants did not have blood samples collected and 2 did not meet inclusion criteria, leaving 53 participants for RNA sequencing analysis. Illnesses were adjudicated as 20 severe and 33 non-severe (14 mild and 19 moderate). This categorization was consistent with the severity separation in the NEWS (Supplemental Figure S1). Two severely ill participants received one dose of Remdesivir prior to blood collection. No subject received steroids or any other experimental COVID-19 treatment prior to enrollment. Five hospitalized participants had rapidly progressive hypoxemia and hemodynamic instability after enrollment and required transfer to intensive care, and 3 subsequently were mechanically ventilated. No mildly ill outpatient illnesses progressed in severity to require medical attention. There was insufficient evidence of any difference in demographic characteristics or underlying conditions by disease severity, except for race and time from disease onset (Table 1): white non-Hispanic comprised 93% mild vs 50-58% moderate-severe (p=0.02), and median time from symptom onset to enrollment was 4 days among mild, 9 days among moderate, and 6.5 days among severe (p=0.047, due to heterogeneity among non-severe). The median age of participants was 62 years with 53% of them being male. As expected, dyspnea, hypoxemia, the presence of infiltrates, and use of supplemental oxygen were more common in moderate and severe, compared to mild illness. All severely ill patients required intensive care; 15 were enrolled in the ICU and 5 were moved to ICU within 48 hours of blood sampling. All severely ill participants required supplemental oxygen; 12 (60%) were mechanically ventilated, one was supported with ECMO and survived, 13 (65%) required vasopressor support and one subject died. Median NEWS were different between the 3 groups (Figure S1). Inflammatory markers were not available for most outpatients but were notably elevated in those hospitalized with moderate to severe disease. (Table 1) Blood gene expression profiling from SARS-CoV-2 positive cases (n=53) was completed by standard mRNA sequencing (RNAseq) of globin mRNA-reduced RNA isolated from whole blood at the time of recruitment. On average 58□±□6 million reads were generated from each of the cDNA libraries, with a mapping rate of 94.2□±□0.6% and transcriptome coverage of 41.3□±□1.3% (Supplemental Figure S2). Exploratory Principal Components Analysis suggested similar patterns of gene expression might be shared by participants with mild and moderate illness, but appeared distinct from those with severe illness (Figure 1A). Statistical analysis for differential gene expression confirmed significant differences when comparing mild vs severe, and moderate vs severe, but not mild vs moderate COVID (Figure 1B).

**Table 1.**
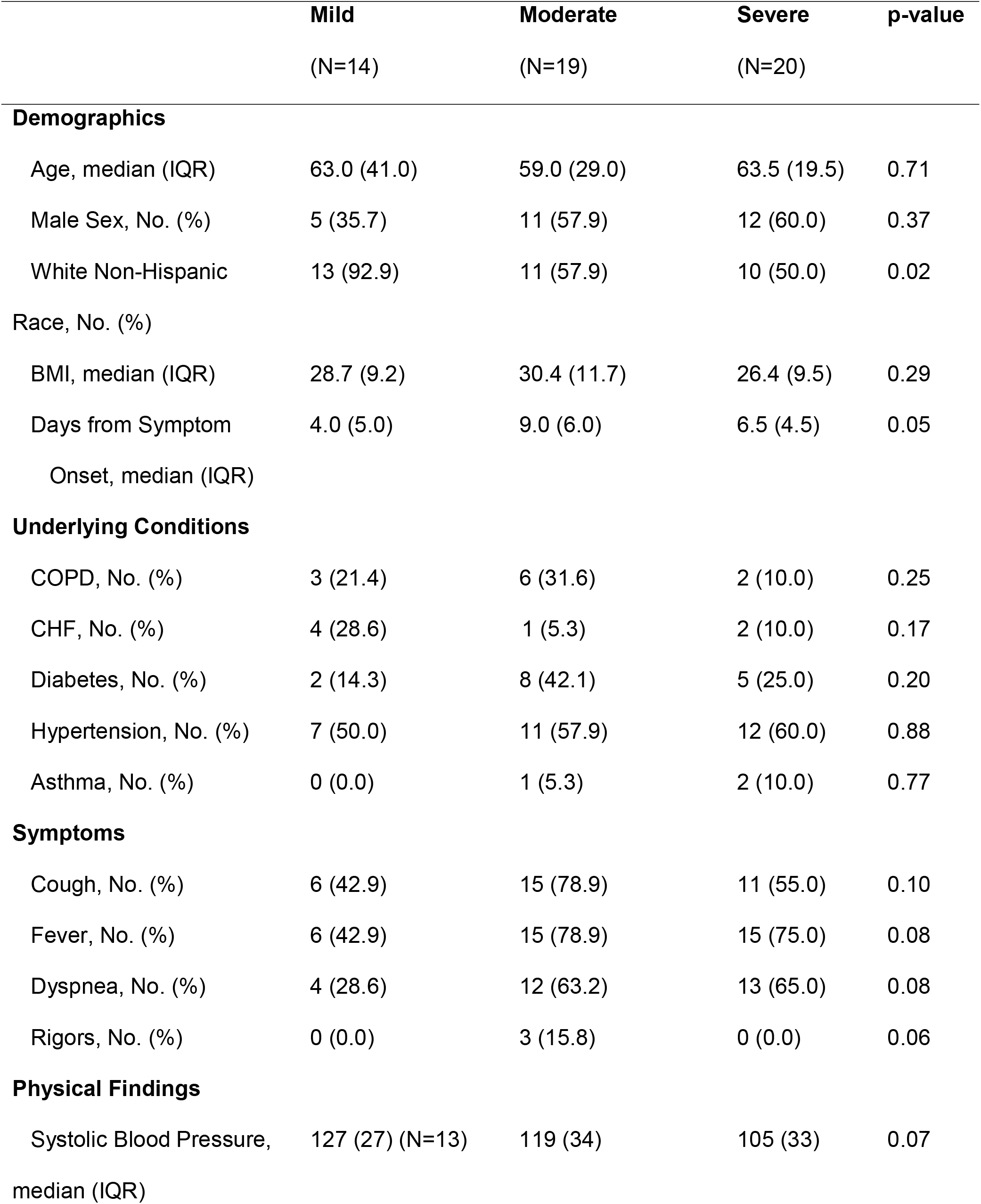
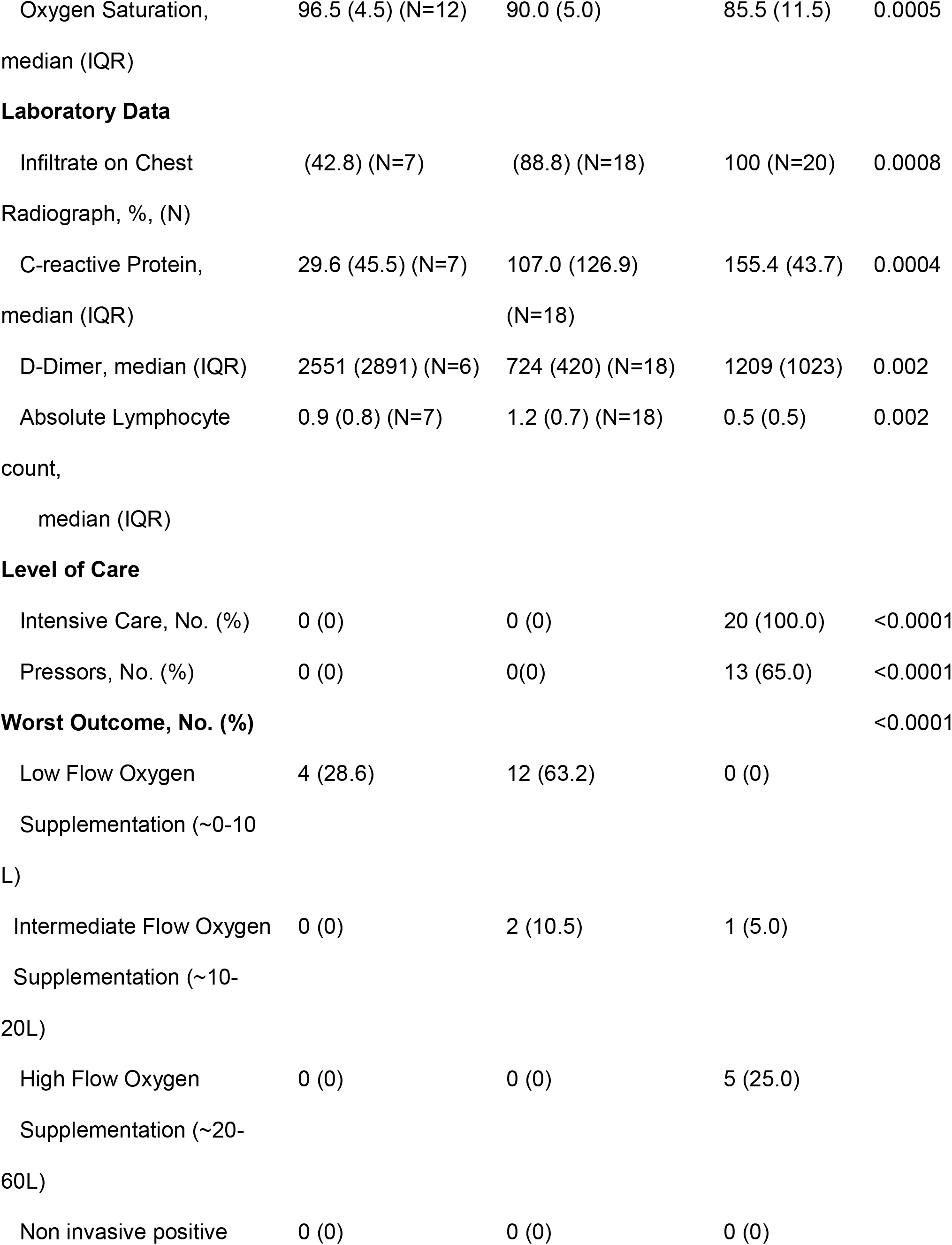
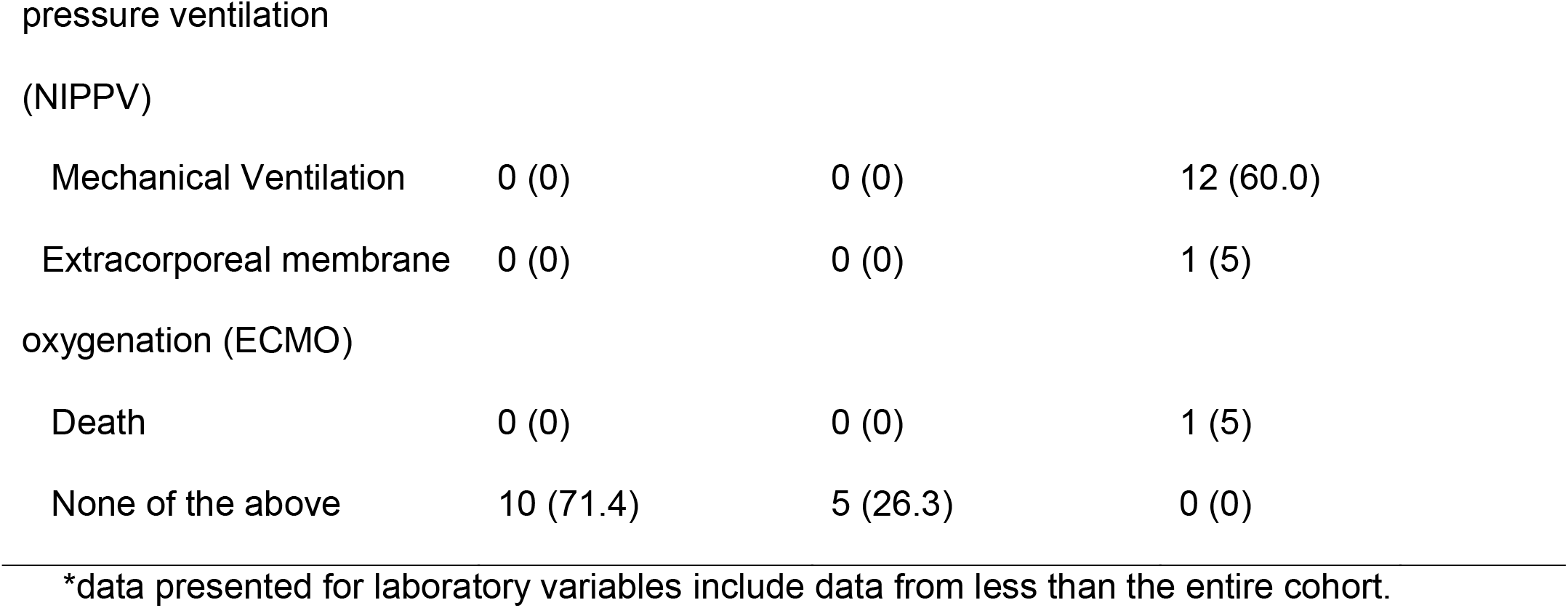
Clinical Variables.

**Figure 1.**
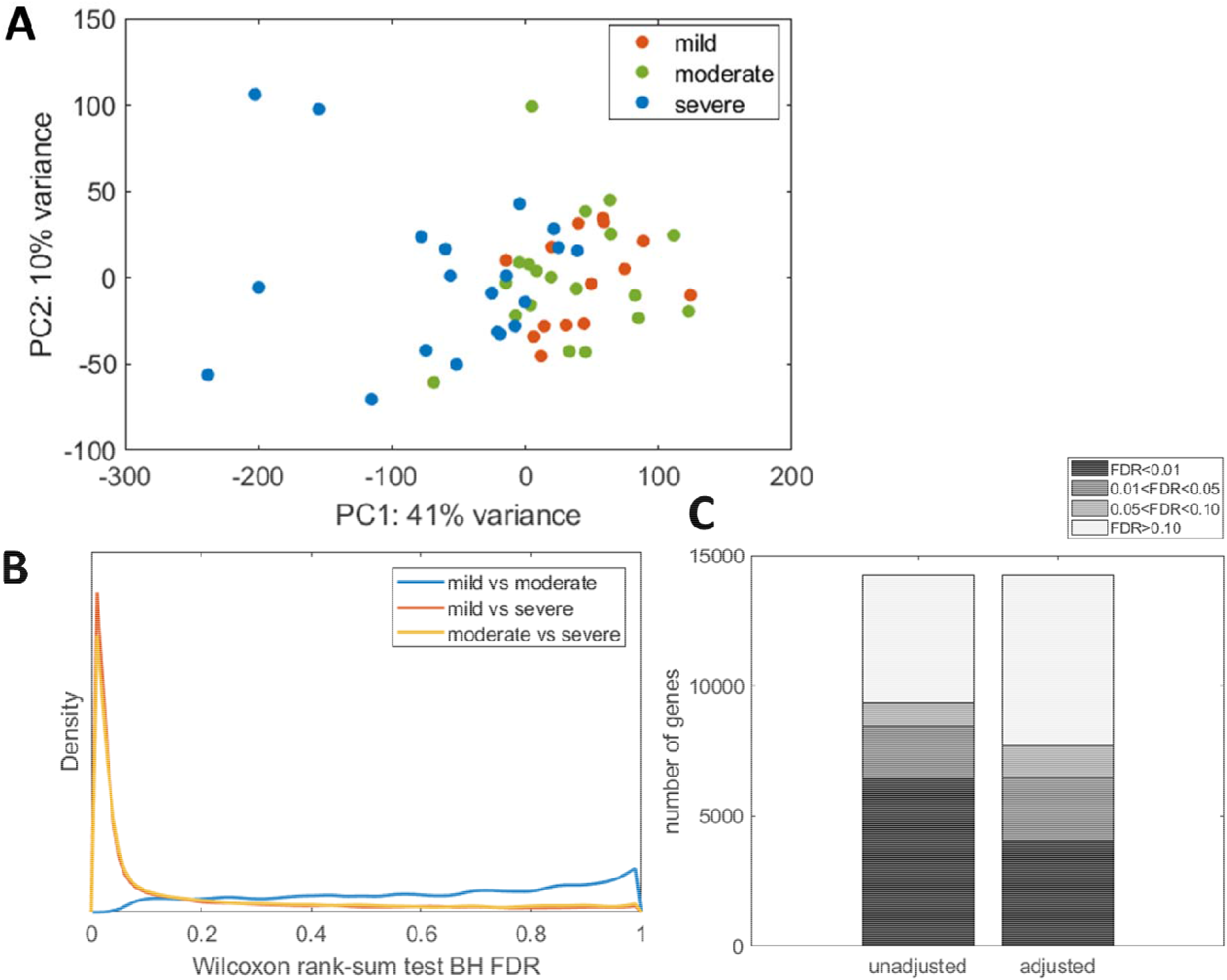
Analyses of signal levels in our dataset which was used as the training data. A. Principal Components Analysis (PCA) plot for Z-score standardized CPM-normalized gene expression, indexed by COVID severity. B. Estimated densities of False Discovery Rates (FDR) for comparisons of COVID severity levels based on nonparametric Wilcoxon tests of CPM-normalized gene expression levels. C. Numbers of differentially expressed genes, by FDR level, based on semiparametric Cox proportional hazards models for gene expression as a function of severe vs non-severe COVID, with and without adjustmentfor pre-specified covariates: race, sex, BMI, days since symptom onset, and library size.

We next tested for differences in gene expression when comparing participants with severe (n=20) vs non-severe illness (n=33), pooling the 14 mild and 19 moderate cases. We tested for differential gene expression without (univariate) and with adjustment for variables potentially associated with severe outcome (race, sex, BMI), the number of days since onset of symptoms, and library size (Figure 1C and Supplemental Table 1). These analyses identified 6483 (46% of tested) and 8435 (59% of tested) differentially expressed genes, with and without multivariate adjustment, respectively.

We performed ontology analysis for the 6483 genes identified as differentially expressed in severe COVID illness, focusing on the fully adjusted analysis (Figure 2). This analysis identified 74 pathways over-represented by genes (n=936) significantly upregulated in severe COVID, and 25 pathways over-represented by genes (n=5547) significantly downregulated (Figure 2 and Supplemental Table S2). Activated pathways included a number associated with infectious diseases as well as TNFα and NFkB signaling. Notably, there was also evidence for significant upregulation of genes associated with platelet activation and coagulation. Among pathways associated with downregulated genes in severe COVID were multiple pathways involved in general host RNA metabolism as well as multiple pathways specifically associated with T cell regulation, including Th2 and Th17 differentiation. The most significantly downregulated pathway was associated with HSV1 infection.

**Figure 2.**
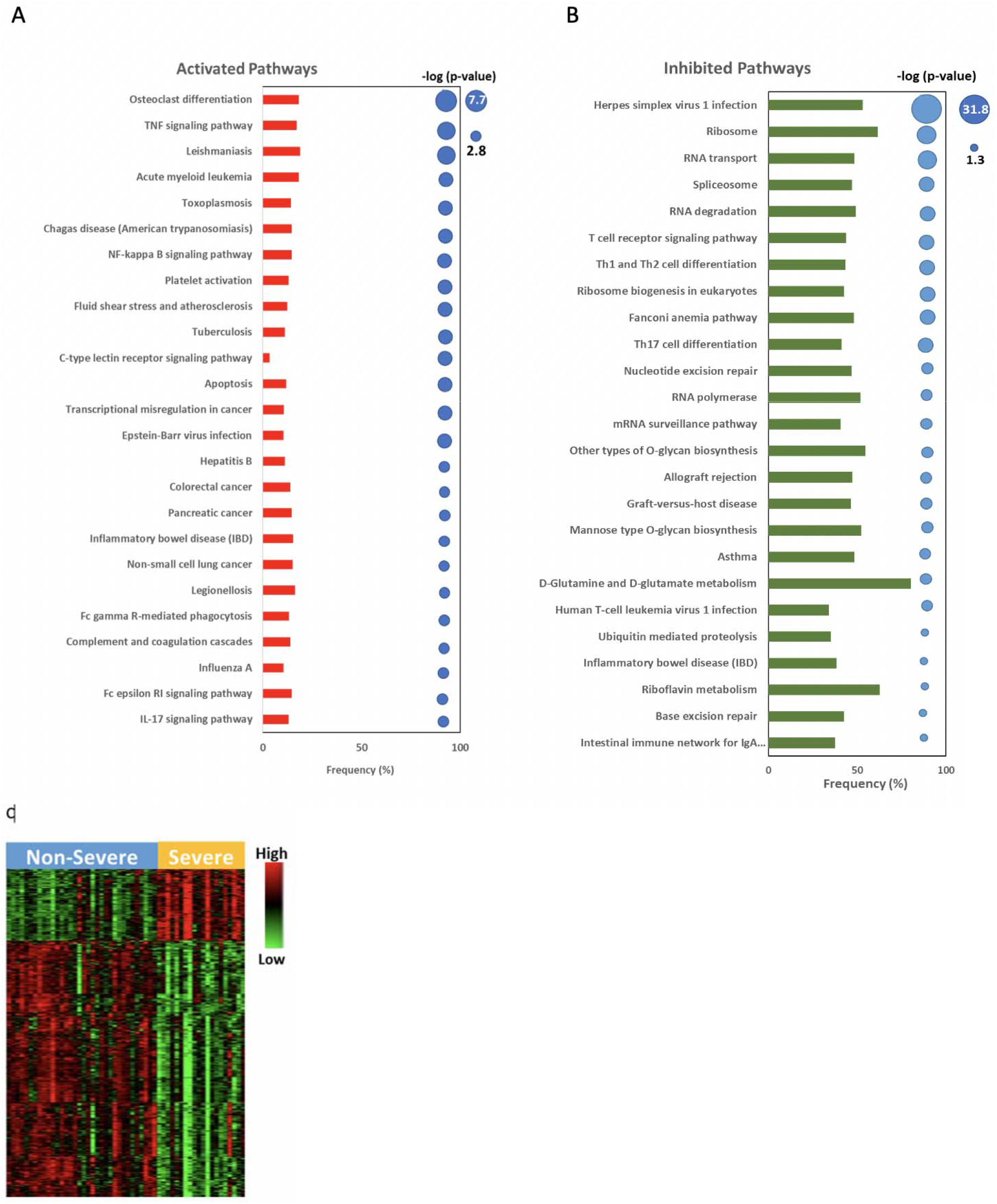
Biological interpretation of gene expression patterns in severe versus non-severe COVID. **(A, B)** Pathway analysis of genes differentially expressed between severe and non-severe COVID-19 participants. Genes identified as overexpressed (n=936), and underexpressed (n=5547) in severe cases of COVID-19, when compared to non-severe cases, were used for pathway analysis using ENRICHR. With 936 genes overexpressed in severe COVID-19 cases, ENRICHR (through KEGG Pathways database) identified 74 pathways associated with COVID-19 severity, while with 5547 genes underexpressed in severe COVID-19, it identified 25 pathways. Shown here are the top 25 significant pathways (p<0.05) associated with upregulated (A) and downregulated genes (B). The bar size represents the frequency of the pathway genes differentially expressed in severe COVID-19; red indicates upregulation, and green indicates downregulation. The size of the dots are proportional to -log(p-value). Larger dots represent lower p-values. **(C)** Differential expression analysis of severe and non-severe COVID-19 participants identified 6483 genes as significantly different. Shown here is a heatmap of the top 425 differentially expressed genes, where the rows indicate genes and columns indicate participants. High expression is shown in red, and low expression in green.

Given the substantial number of differentially expressed genes when comparing severe vs non-severe COVID, we investigated the ability of gene expression patterns to discriminate severe illness. Gene-specific thresholds for univariate AUC and magnitude change were chosen via the cross-validation procedure and used to produce an 18 gene weighted gene expression risk score (WGERS) for severe illness. Nested cross-validation was used to estimate performance via the stratified AUC (CV-AUC=0.98). The pooled CV-AUC of 0.93 corresponds with a cross-validated ROC curve to graphically summarize performance (Figure 3A). The pooled CV-ROC curve also was used to select a risk score threshold (−1.04) with 95% sensitivity and 88% specificity, which corresponded with apparent (non-cross-validated) sensitivity of 100%, specificity of 85%, and error rate of 9% (5/53), represented via the WGERS distributions for the training data (Figure 4A). All 5 misclassified participants had moderate illness (Figure 4B).

**Figure 3.**
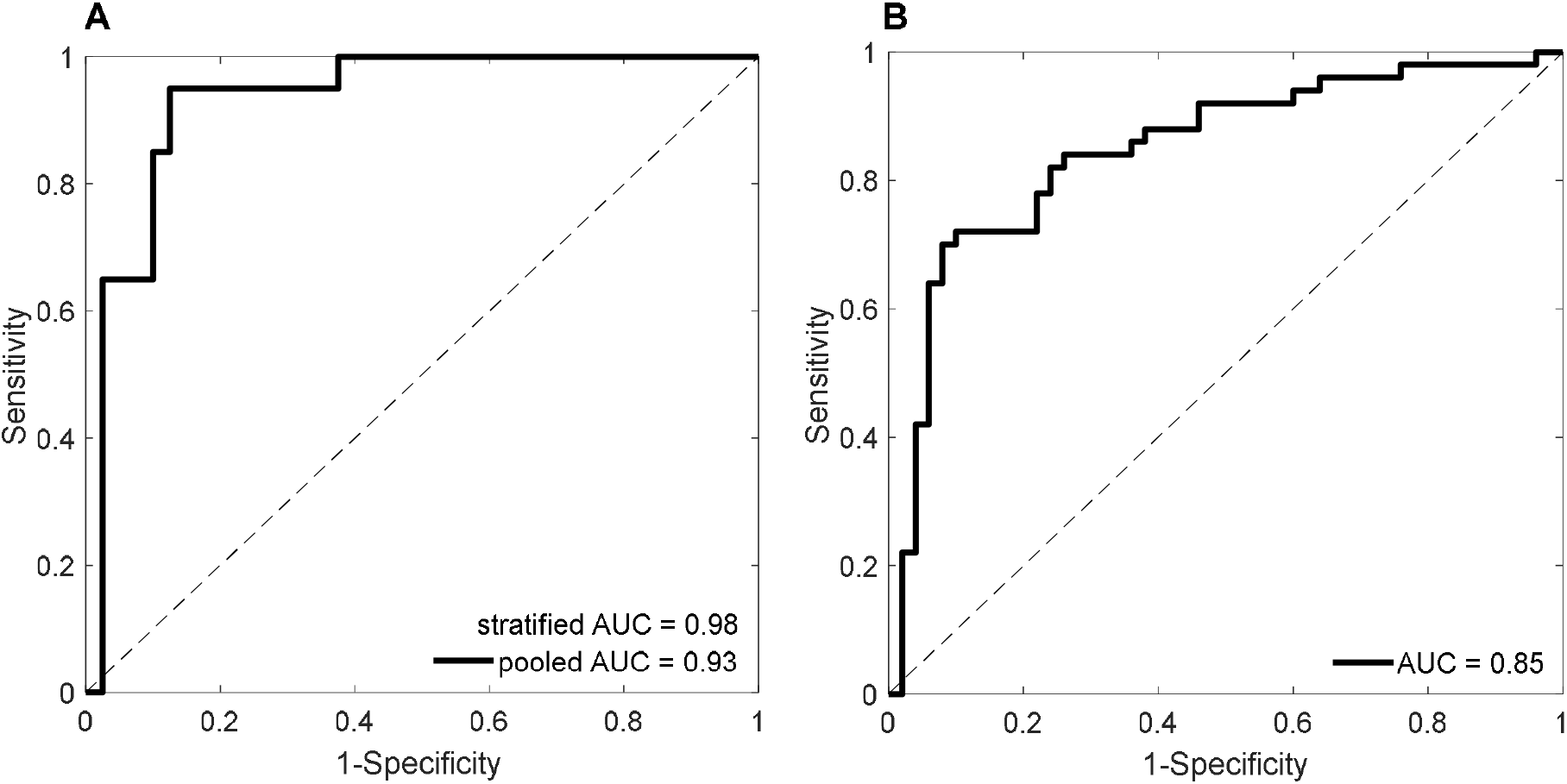
Internally cross-validated and externally validated Receiver Operating Characteristic (ROC) curves for the Weighted Gene Expression Risk Score (WGERS). A. Cross-Validated (CV) ROC for severe vs non-severe COVID in the Training Data. Pooled AUC corresponds with the plot (95% sensitivity and 88% specificity at a risk score threshold of -1.04), which necessarily compares risk scores both within and across CV folds (each with a unique fitted model). Stratified AUC corresponds with only comparing risk scores within each fold of 20-fold CV (where each model is fixed), but there exists no single corresponding ROC curve for this more commonly reported and preferable metric. B. ROC curve for ICU vs non-ICU in the Validation Data. A WGERS threshold of 1.77 yielded 84% sensitivity and 74% specificity.

**Figure 4.**
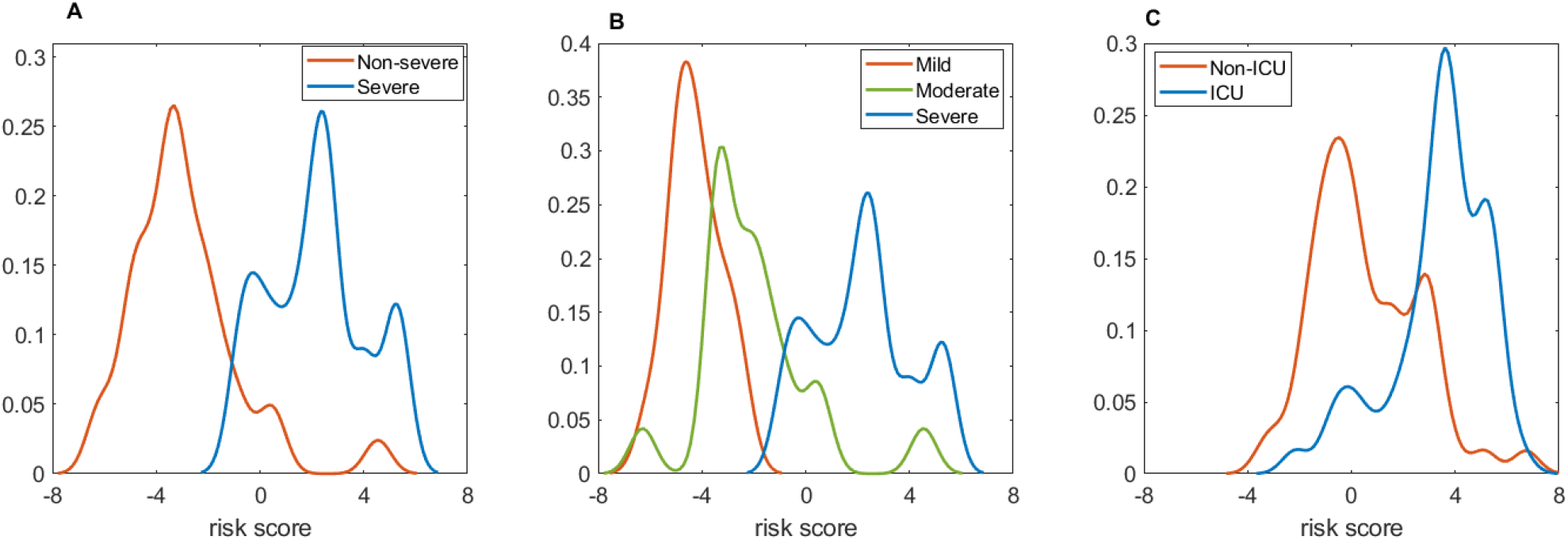
Risk score distributions in the Training and Validation Data. A. Density of risk scores by non-severe (mild or moderate) vs severe COVID in the Training Data. Apparent (non-cross-validated) sensitivity = 100% (20/20 severe) and specificity = 85% (28/33 non-severe) at the CV-optimal risk score threshold of -1.04. B. Density of risk scores by COVID severity (mild, moderate, or severe) in the Training Data. Although the statistical learner was blinded to any distinction between mild and moderate COVID severity, risk scores for moderate COVID participants fell between those of mild and severe COVID participants, and all 5 misclassified participants had moderate COVID. C. Density of risk scores by ICU vs non-ICU in the Validation Data. Sensitivity = 84% (42/50 ICU) and specificity = 74% (37/50 non-ICU) at a risk score threshold of 1.77.

We next identified an independent validation data set describing peripheral blood-based gene expression profiling of COVID subjects who were either admitted (n=50) or not admitted (n=50) to the ICU due to the severity of their acute illness [14]. Our 18 gene WGERS discriminated between ICU and non-ICU patients with an AUC of 0.85, and thresholding at 1.77 yielded 84% sensitivity and 74% specificity (Figures 3B and 4C). Furthermore, all 18 genes selected in the training data were differentially expressed (FDR < 0.01) in the validation data (Supplemental Table S3).

## Discussion

SARS-CoV-2 infection causes a wide spectrum of disease ranging from minimal, often asymptomatic, respiratory illness to severe pneumonia with multisystem failure and death. Although measurements of inflammatory markers such as C-reactive protein and serum IL-6 levels are often associated with worse disease, their use to predict poor outcomes is imperfect [15-17]. Viral characteristics, such as shedding kinetics or gene sequence variation, are not reliable predictors of clinical outcome [18, 19]. Genome-wide expression profiling, a powerful and unbiased tool, can be used for multiple purposes such as relating activation or suppression of molecular pathways to clinical manifestations of disease, identification of biomarkers that may allow individual prediction of disease severity, and identification of novel gene targets for therapeutic intervention. Early predictors to identify patients that will decompensate following SARS-CoV-2 infection would be highly impactful.

In our study of 53 SARS-CoV-2 infected adults with illness ranging from very mild upper respiratory infection to acute respiratory failure, we identified >6,000 differentially expressed genes (DEGs) (FDR < 0.05) between severe and non-severe illness. The vast majority (85%) of DEGs were under-expressed, most notably with a marked effect on lymphocytes and altered function [20, 21]. Pathway analysis revealed inhibition of Th1, Th2 and Th17 cell differentiation, as well as inhibition of the T cell receptor signaling pathway. These effects are likely related to the marked lymphopenia and poor adaptive immune response in persons with severe SARS-CoV-2 infection [22]. Also notable in severe illness is the inhibition of the mRNA surveillance pathways that include the nonsense-mediated mRNA decay pathway which can degrade viral mRNA. Using a model coronavirus, murine hepatitis virus, Wada and colleagues showed viral transcription is enhanced by blocking this host cell pathway, demonstrated to be mediated by the viral nucleocapsid protein [23].

Several activated pathways we identified in our studies are worth comment, given what is already known about SARS-CoV-2 and COVID-19. Activation of the NF-kappa B and TNF signaling pathways in a setting of heightened inflammatory process is not surprising. Activation of the platelet, complement, and coagulation cascade pathways are also expected, given the characteristic hypercoagulable state that has been observed in severe illness [24]. Thrombocytopenia and activated platelets are associated with the high incidence of venous and arterial clotting, while elevated levels of serum D-dimer, a fibrinogen degradation product, and increased INR are all features of severe COVID-19 [25]. It is interesting that the infection-related pathways most significantly activated include those principally associated with intracellular bacterial (legionella, mycobacterial) and parasitic (toxoplasma, leishmania and trypanosome [Chagas]) infections. These infections are associated with marked activation of macrophages, and thus may be consistent with activation of the osteoclast differentiation pathway, as osteoclasts and macrophages have many similarities [26, 27].

Our findings are generally consistent with the limited data currently available in the literature on COVID-19 and gene expression [14, 28-31]. Specifically, Overmyer et al reported gene expression and metabolomic data from 128 COVID infected and non COVID infected persons, where 219 molecular features with high significance to COVID-19 status and severity were discovered. [14] A number of these involved complement activation, dysregulated lipid transport, and neutrophil activation. Additionally, our data is supported by the findings noted by Ouyang et al in which Th17 and T cell activation and differentiation were markedly downregulated in severe disease [32].

There were some novel pathways that demonstrated upregulation in our studies. A number of malignancy related pathways were upregulated (acute myelogenous leukemia (AML), colorectal, pancreatic and Non-small cell cancer) in the severe COVID patients. Similarly, in a study by Kwan et al comparing gene expression in 45 COVID-19 cases to healthy controls, 135 genes were found to be differentially expressed, with enrichment for several cancer pathways including viral carcinogenesis and AML [29]. Identification of mutations in cell signaling, proliferation genes, and kinases such as AP2-associated protein kinase 1 (AAK1) have led to targeted treatment options for cancer patients. Baricitinib, a repurposed rheumatoid arthritis drug that interferes with the Janus Kinase (JAK) pathway, was demonstrated to have efficacy when combined with Remdesivir in the treatment of severe COVID [33]. Baricitinib shows high affinity for AP2 associated protein kinase 1 binding, potentially demonstrating some overlap in perturbations of cell signaling pathways in malignancy and COVID-19. Further investigation of gene expression pathways differentially expressed in severely ill patients may provide clues to new therapeutic targets.

Although our study was not designed to identify and validate early predictors of severe disease, the data do offer a first step. Using gene expression data we were able develop and validate an 18 gene signature for severe disease –fully concordant with requiring ICU– with 85% AUC, 84% sensitivity, and 74% specificity in an independent validation data set. In a recent paper Guardela et al assessed the utility of blood transcript levels of 50 genes known to predict mortality in Idiopathic Pulmonary Fibrosis patients to classify illness severity in COVID-19 [31]. A discovery cohort of eight subjects was used, and then validated using a publicly available data set of 128 subjects [14]. The gene expression risk profile discriminated ICU admission, need for mechanical ventilation, and in-hospital mortality with an AUC of 77%, 75%, and 74%, respectively (p < 0.001) in a COVID-19 validation cohort.

Our current study has several limitations which are worth noting, including its relatively small sample size, the non-standardized interval between symptom onset and sample collection, and blood collection at one time point. The complexity of the clinical data among hospitalized participants (i.e. admissions only for isolation, persons with chronic oxygen requirements, COVID testing for procedures) made objective criteria to distinguish mild from moderate disease difficult, necessitating the need for clinical adjudication. Lastly, certain laboratory studies were not available for all subjects.

## Conclusions

In summary, we found a large number of differentially expressed genes in the peripheral blood that distinguished those with severe COVID-19 illness from those with mild or moderate disease. These data could be used to identify potential targets for interventions, as well as to develop predictors of disease severity. Future prospective studies are needed to follow mild to moderately ill patients over time and evaluate whether any of the discriminatory genes identified are affected at early stages and can serve as predicators of severity. If so, individuals with high risk gene profiles might be hospitalized for observation, moved to a more closely monitored setting while hospitalized, or targeted for early interventions such as monoclonal antibody treatments.

## Acknowledgements

We acknowledge the University of Rochester Genomics Research Center for completing RNA isolation and sequencing. Christopher Slaunwhite, Mary Anne Formica, and Michael Peasley provided technical support. Jeanne Holden-Wiltse and Jeffrey Williams assisted with data management.

## Author Contributions

DRP, EEW, ARF and TJM conceptualized the study. DPC, ARB, EEW and ARF recruited patients, adjudicated clinical severity and collected samples. DRP, AMB, SB, AMC, and TJM analyzed the data. DRP, AMB, SB, ARB, DPC, EEW, TJM and ARF interpreted the results. All authors contributed to writing of the manuscript. All authors reviewed, edited and approved the manuscript for submission.

The manuscript represents original work that is not currently under consideration elsewhere.

## Data Availability

Raw and processed data from the study are currently in process of being submitted to DBGAP

