## Supplemental Tables and Figures for "Gene Expression Risk Scores for COVID-19 Illness Severity"

University of Rochester Medical Center

601 Elmwood Ave, Box 850

Rochester, NY 14642, USA.

Ann R. Falsey, M.D.

Infectious Disease Unit

Rochester General Hospital

1425 Portland Avenue

Rochester, NY, 14621 USA

Running Title: Risk Scores for COVID-19 Severity

3 Supplementary Tables

2 Supplementary Figures

Supplemental Figure S1. Distribution of National Early Warning Score (NEWS) at the time of RNAseq blood collection by COVID severity. Box plots display median NEWS, plus upper and lower quartiles, with whiskers depicting the range.

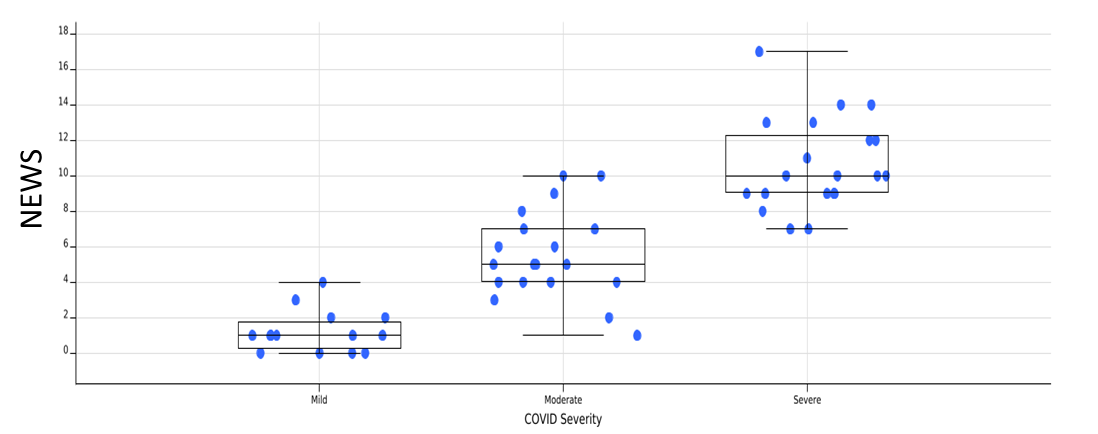

Supplemental Figure S2: Transcriptomics data quality metrics by COVID severity.

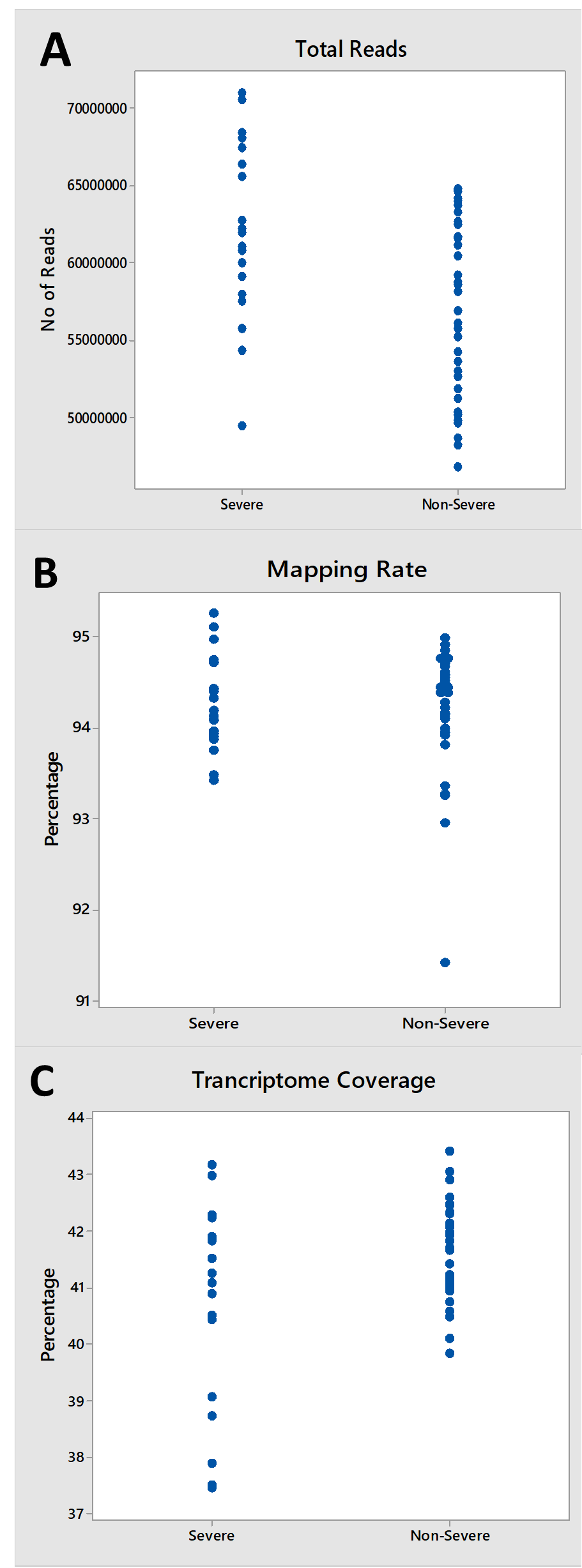

(A) total number of input reads, (B) percentage of input reads mapped to the genome, and (C) the proportion of the genome for which transcripts were detected, grouped by severity level.

Supplemental Table S1. Numbers of differentially expressed genes for severe vs non-severe COVID, by FDR level, with and without adjustment for pre-specified covariates.

| **Adjustment** | **Min(FDR)** | **# Genes FDR < 0.01** | **# Genes FDR < 0.05** | **# Genes FDR < 0.10** |
| --- | --- | --- | --- | --- |
| *None* | 2.48E-7 | 6444 | 8435 | 9364 |
| *Race, sex, BMI, days since symptom onset, and library size* | 1.04E-5 | 4060 | 6483 | 7721 |

Supplemental Table S2: Pathways associated with COVID-19 Severity

| **Pathways** | **Proportion*** | **P-value** | **Genes** |
| --- | --- | --- | --- |
| *Activated* | *Pathways* |  |  |
| Osteoclast differentiation | 23/127 | 2.18E-08 | JUN;SPI1;TGFB1;IL1R1;IFNGR1;IFNGR2;CYBA;PIK3CB;MAPK14;LILRB3;TNFRSF1A;RELB;LILRA5;NFKBIA;SOCS3;FCGR2A;SIRPA;GRB2;PPARG;IKBKG;JUNB;MAP2K6;MAPK3 |
| TNF signaling pathway | 19/110 | 7.73E-07 | MAP2K3;JUN;CEBPB;TNFAIP3;PIK3CB;CFLAR;MAPK14;MMP9;ICAM1;TNFRSF1A;NFKBIA;SOCS3;BCL3;FADD;IKBKG;JUNB;IL18R1;MAP2K6;MAPK3 |
| Leishmaniasis | 14/74 | 7.26E-06 | IL10;JUN;ITGAM;CR1;TGFB1;IFNGR1;IFNGR2;CYBA;MAPK14;NFKBIA;FCGR2A;TLR4;MYD88;MAPK3 |
| Acute myeloid leukemia | 12/66 | 4.90E-05 | CCNA1;STAT5B;SPI1;ITGAM;BCL2A1;FLT3;RARA;GRB2;PIK3CB;IKBKG;MPO;MAPK3 |
| Toxoplasmosis | 16/113 | 7.28E-05 | MAP2K3;IL10;TGFB1;IFNGR1;IFNGR2;LAMC1;MAPK14;TNFRSF1A;NFKBIA;CASP9;ALOX5;IKBKG;TLR4;MYD88;MAP2K6;MAPK3 |
| Chagas disease (American trypanosomiasis) | 15/103 | 8.75E-05 | IL10;JUN;TGFB1;IFNGR1;IFNGR2;PIK3CB;CFLAR;MAPK14;TNFRSF1A;NFKBIA;FADD;IKBKG;TLR4;MYD88;MAPK3 |
| NF-kappa B signaling pathway | 14/95 | 1.31E-04 | LYN;BCL2A1;GADD45B;TNFSF14;IL1R1;TNFAIP3;CFLAR;ICAM1;TNFRSF1A;RELB;NFKBIA;IKBKG;TLR4;MYD88 |
| Platelet activation | 16/124 | 2.21E-04 | VASP;LYN;ITGA2B;ADCY4;ADCY3;GP1BA;PIK3CB;MAPK14;PRKCZ;APBB1IP;GP9;FCGR2A;P2RX1;TBXAS1;FERMT3;MAPK3 |
| Fluid shear stress and atherosclerosis | 17/139 | 2.73E-04 | JUN;IL1R1;IL1R2;ITGA2B;CYBA;TXN;PIK3CB;MAPK14;PRKCZ;MMP9;ICAM1;TNFRSF1A;THBD;RAC2;SDC1;IKBKG;MAP2K6 |
| Tuberculosis | 20/179 | 2.82E-04 | IL10;PLK3;CEBPB;TGFB1;ITGAM;CR1;IFNGR1;IFNGR2;MAPK14;TNFRSF1A;CASP9;FCGR2A;IRAK2;ITGAX;FADD;CTSD;TLR4;MYD88;RAB7A;MAPK3 |
| C-type lectin receptor signaling pathway | 14/104 | 3.45E-04 | IL10;PLK3;JUN;PRKCD;PIK3CB;MAPK14;RELB;NFKBIA;CLEC4D;MAPKAPK2;BCL3;IKBKG;CLEC1B;MAPK3 |
| Apoptosis | 17/143 | 3.83E-04 | JUN;GADD45B;BCL2A1;GADD45A;HTRA2;PIK3CB;CFLAR;TUBA4A;GADD45G;LMNB1;TNFRSF1A;NFKBIA;CASP9;FADD;IKBKG;CTSD;MAPK3 |
| Transcriptional misregulation in cancer | 20/186 | 4.67E-04 | CEBPB;SPI1;ITGAM;GADD45B;H3F3B;HPGD;BCL2A1;GADD45A;FLT3;IL1R2;TFE3;MLLT1;MPO;MMP9;GADD45G;CCNA1;BCL6;RARA;PPARG;ELANE |
| Epstein-Barr virus infection | 21/201 | 5.01E-04 | LYN;MAP2K3;JUN;GADD45B;GADD45A;TNFAIP3;PIK3CB;MAPK14;ICAM1;HLA-E;GADD45G;RELB;NFKBIA;CASP9;CCNA1;FADD;IKBKG;CD58;JAK3;MYD88;MAP2K6 |
| Hepatitis B | 18/163 | 6.41E-04 | MAP2K3;STAT5B;JUN;TGFB1;PIK3CB;MAPK14;MMP9;NFKBIA;CASP9;CCNA1;GRB2;FADD;IKBKG;JAK3;TLR4;MYD88;MAP2K6;MAPK3 |
| Colorectal cancer | 12/86 | 6.46E-04 | CASP9;JUN;RALB;TGFB1;GADD45B;GADD45A;RAC2;TGFA;GRB2;PIK3CB;MAPK3;GADD45G |
| Pancreatic cancer | 11/75 | 7.01E-04 | CASP9;RALB;TGFB1;GADD45B;GADD45A;RAC2;TGFA;PIK3CB;IKBKG;MAPK3;GADD45G |
| Inflammatory bowel disease (IBD) | 10/65 | 8.28E-04 | IL10;JUN;IL18RAP;IL4R;TGFB1;IFNGR1;IFNGR2;TLR5;TLR4;IL18R1 |
| Non-small cell lung cancer | 10/66 | 9.36E-04 | CASP9;STAT5B;GADD45B;GADD45A;TGFA;GRB2;PIK3CB;JAK3;MAPK3;GADD45G |
| Legionellosis | 9/55 | 9.47E-04 | CASP9;NFKBIA;RAB1A;ITGAM;CR1;NLRC4;TLR5;TLR4;MYD88 |
| Fc gamma R-mediated phagocytosis | 12/91 | 0.0010768 | LYN;VASP;HCK;GSN;FCGR2A;MYO10;LIMK2;PRKCD;RAC2;AMPH;PIK3CB;MAPK3 |
| Complement and coagulation cascades | 11/79 | 0.0010909 | THBD;SERPINA1;ITGAM;CR1;C1R;C5AR1;ITGAX;CD59;TFPI;CLU;CD55 |
| Influenza A | 18/171 | 0.0011236 | MAP2K3;JUN;IFNGR1;IFNGR2;FURIN;PIK3CB;MAPK14;ICAM1;TNFRSF1A;NFKBIA;CASP9;DNAJC3;SOCS3;TLR4;MYD88;AGFG1;MAP2K6;MAPK3 |
| Fc epsilon RI signaling pathway | 10/68 | 0.0011854 | MAP2K3;LYN;ALOX5;ALOX5AP;RAC2;GRB2;PIK3CB;MAPK14;MAPK3;MAP2K6 |
| IL-17 signaling pathway | 12/93 | 0.0013057 | NFKBIA;CEBPB;JUN;TNFAIP3;IKBKG;FADD;MAPK14;MMP9;S100A9;S100A8;IL17RA;MAPK3 |
| Th17 cell differentiation | 13/107 | 0.0014679 | STAT5B;JUN;IL4R;TGFB1;IL1R1;IFNGR1;IFNGR2;MAPK14;NFKBIA;RARA;IKBKG;JAK3;MAPK3 |
| Chemokine signaling pathway | 19/190 | 0.0015278 | LYN;STAT5B;XCR1;PRKCD;PXN;ADCY4;ADCY3;PIK3CB;PRKCZ;NFKBIA;FGR;HCK;GNB2;GRK6;RAC2;GRB2;IKBKG;JAK3;MAPK3 |
| Amoebiasis | 12/96 | 0.0017234 | IL10;SERPINB10;SERPINB1;ITGAM;TGFB1;IL1R1;ARG1;IL1R2;PIK3CB;LAMC1;TLR4;RAB7A |
| Malaria | 8/49 | 0.0018272 | IL10;CR1;TGFB1;SDC1;TLR4;MYD88;ICAM1;THBS3 |
| Hematopoietic cell lineage | 12/97 | 0.0018848 | GP9;CSF3R;IL4R;ITGAM;CR1;IL1R1;FLT3;IL1R2;ITGA2B;GP1BA;CD59;CD55 |
| Salmonella infection | 11/86 | 0.0021968 | JUN;IFNGR1;IFNGR2;RHOG;NLRC4;TLR5;MAPK14;TLR4;MYD88;RAB7A;MAPK3 |
| HIF-1 signaling pathway | 12/100 | 0.0024451 | HK3;LDHA;PFKFB3;IFNGR1;IFNGR2;MKNK1;PGK1;PIK3CB;ALDOA;TLR4;GAPDH;MAPK3 |
| Chronic myeloid leukemia | 10/76 | 0.0027825 | NFKBIA;STAT5B;TGFB1;GADD45B;GADD45A;GRB2;PIK3CB;IKBKG;MAPK3;GADD45G |
| Kaposi sarcoma-associated herpesvirus infection | 18/186 | 0.0028722 | LYN;JUN;IFNGR1;PIK3CB;MAPK14;ICAM1;HLA-E;TNFRSF1A;NFKBIA;CASP9;HCK;ZFP36;GNB2;MAPKAPK2;FADD;IKBKG;MAP2K6;MAPK3 |
| Toll-like receptor signaling pathway | 12/104 | 0.003396 | MAP2K3;NFKBIA;JUN;FADD;PIK3CB;IKBKG;TLR5;MAPK14;TLR4;MYD88;MAP2K6;MAPK3 |
| Human T-cell leukemia virus 1 infection | 20/219 | 0.0034063 | STAT5B;JUN;SPI1;TGFB1;IL1R1;IL1R2;ADCY4;ADCY3;PIK3CB;ETS2;ICAM1;HLA-E;RELB;TNFRSF1A;NFKBIA;CCNA1;ZFP36;IKBKG;JAK3;MAPK3 |
| Insulin signaling pathway | 14/137 | 0.0049811 | PYGM;PIK3CB;PYGL;PHKA2;PRKCZ;SOCS3;HK3;MKNK1;PPP1R3D;FLOT1;FLOT2;GRB2;SH2B2;MAPK3 |
| Human immunodeficiency virus 1 infection | 19/212 | 0.0051958 | MAP2K3;JUN;LIMK2;PXN;PIK3CB;MAPK14;HLA-E;TNFRSF1A;NFKBIA;CASP9;GNB2;RAC2;FADD;IKBKG;TLR4;AP1M2;MYD88;MAP2K6;MAPK3 |
| Type II diabetes mellitus | 7/46 | 0.0052167 | HK3;SOCS3;PRKCD;PIK3CB;PRKCZ;CACNA1E;MAPK3 |
| B cell receptor signaling pathway | 9/71 | 0.0057126 | LYN;NFKBIA;JUN;RAC2;GRB2;PIK3CB;IKBKG;LILRB3;MAPK3 |
| Pathways in cancer | 38/530 | 0.0061992 | SPI1;CSF3R;RALB;FLT3;EPAS1;ITGA2B;ADCY4;ADCY3;TGFA;PIK3CB;LAMC1;RASGRP4;CASP9;FRAT1;WNT11;RAC2;IKBKG;FADD;JAK3;MAPK3;STAT5B;JUN;IL4R;TGFB1;GADD45B;IFNGR1;GADD45A;DAPK2;IFNGR2;TXNRD1;MMP9;GADD45G;NFKBIA;CCNA1;GNB2;RARA;PPARG;GRB2 |
| MAPK signaling pathway | 24/295 | 0.0062506 | MAP2K3;JUN;TGFB1;GADD45B;IL1R1;GADD45A;FLT3;TGFA;MAPK14;CACNA1E;RASGRP4;GADD45G;RELB;TNFRSF1A;MKNK1;RPS6KA1;MAPKAPK2;RAC2;MAP3K20;GRB2;IKBKG;MYD88;MAP2K6;MAPK3 |
| Viral carcinogenesis | 18/201 | 0.0064664 | LYN;STAT5B;JUN;GSN;HPN;PXN;HIST1H2BK;PIK3CB;HLA-E;HDAC7;NFKBIA;CCNA1;MAPKAPK2;GRB2;IKBKG;JAK3;YWHAH;MAPK3 |
| Relaxin signaling pathway | 13/130 | 0.0079621 | JUN;TGFB1;ADCY4;ADCY3;PIK3CB;MAPK14;PRKCZ;MMP9;NFKBIA;GNB2;GRB2;INSL3;MAPK3 |
| Necroptosis | 15/162 | 0.009163 | STAT5B;IFNGR1;IFNGR2;TNFAIP3;PYGM;PYGL;CFLAR;TNFRSF1A;SPATA2L;SMPD1;CHMP2A;CHMP3;FADD;JAK3;TLR4 |
| NOD-like receptor signaling pathway | 16/178 | 0.0095486 | JUN;PRKCD;TNFAIP3;CYBA;TXN;NLRC4;MAPK14;NFKBIA;NLRP6;MFN2;FADD;IKBKG;TLR4;MYD88;MCU;MAPK3 |
| Neurotrophin signaling pathway | 12/119 | 0.009894 | NFKBIA;JUN;IRAK2;RPS6KA1;PRKCD;MAPKAPK2;GRB2;IRAK3;PIK3CB;MAPK14;SH2B2;MAPK3 |
| Central carbon metabolism in cancer | 8/65 | 0.0106301 | HK3;SLC7A5;G6PD;LDHA;FLT3;PIK3CB;SLC16A3;MAPK3 |
| Shigellosis | 8/65 | 0.0106301 | NFKBIA;CTTN;RHOG;ELMO2;HCLS1;IKBKG;MAPK14;MAPK3 |
| Th1 and Th2 cell differentiation | 10/92 | 0.0107926 | NFKBIA;STAT5B;JUN;IL4R;IFNGR1;IFNGR2;IKBKG;MAPK14;JAK3;MAPK3 |
| Pentose phosphate pathway | 5/30 | 0.0119451 | G6PD;H6PD;IDNK;PGD;ALDOA |
| Staphylococcus aureus infection | 8/68 | 0.013783 | IL10;ITGAM;FCGR2A;C1R;C5AR1;FPR1;FCAR;ICAM1 |
| Renal cell carcinoma | 8/69 | 0.0149714 | JUN;TGFB1;EPAS1;TFE3;TGFA;GRB2;PIK3CB;MAPK3 |
| Rap1 signaling pathway | 17/206 | 0.0169878 | VASP;MAP2K3;ITGAM;RALB;ITGA2B;FPR1;ADCY4;ADCY3;PIK3CB;SIPA1L2;MAPK14;PRKCZ;APBB1IP;CNR1;RAC2;MAP2K6;MAPK3 |
| Fructose and mannose metabolism | 5/33 | 0.0177358 | PFKFB2;HK3;PFKFB4;PFKFB3;ALDOA |
| Endometrial cancer | 7/58 | 0.0181132 | CASP9;GADD45B;GADD45A;GRB2;PIK3CB;MAPK3;GADD45G |
| AGE-RAGE signaling pathway in diabetic complications | 10/100 | 0.0186096 | STAT5B;THBD;JUN;TGFB1;PRKCD;PIK3CB;MAPK14;PRKCZ;ICAM1;MAPK3 |
| VEGF signaling pathway | 7/59 | 0.0197487 | CASP9;MAPKAPK2;PXN;RAC2;PIK3CB;MAPK14;MAPK3 |
| FoxO signaling pathway | 12/132 | 0.0210935 | IL10;CDKN2D;PLK3;TGFB1;GADD45B;BCL6;GADD45A;GRB2;PIK3CB;MAPK14;GADD45G;MAPK3 |
| Starch and sucrose metabolism | 5/36 | 0.0251284 | HK3;MGAM;PYGM;PYGL;GYG1 |
| Pertussis | 8/76 | 0.0254466 | IL10;JUN;ITGAM;C1R;MAPK14;TLR4;MYD88;MAPK3 |
| Estrogen signaling pathway | 12/137 | 0.0272677 | JUN;PRKCD;RARA;ADCY4;TGFA;ADCY3;GRB2;PIK3CB;CTSD;MMP9;FKBP5;MAPK3 |
| Thyroid cancer | 5/37 | 0.0279715 | GADD45B;GADD45A;PPARG;MAPK3;GADD45G |
| GnRH signaling pathway | 9/93 | 0.0299471 | MAP2K3;JUN;PRKCD;ADCY4;ADCY3;GRB2;MAPK14;MAPK3;MAP2K6 |
| Small cell lung cancer | 9/93 | 0.0299471 | CASP9;NFKBIA;GADD45B;GADD45A;ITGA2B;PIK3CB;LAMC1;IKBKG;GADD45G |
| Insulin resistance | 10/108 | 0.0299516 | NFKBIA;SOCS3;RPS6KA1;PRKCD;PPP1R3D;PYGM;PIK3CB;PYGL;PRKCZ;TNFRSF1A |
| Human cytomegalovirus infection | 17/225 | 0.0358648 | IL1R1;PXN;ADCY4;ADCY3;PIK3CB;MAPK14;HLA-E;TNFRSF1A;NFKBIA;CASP9;GNB2;RAC2;GRB2;FADD;IKBKG;MAP2K6;MAPK3 |
| Ferroptosis | 5/40 | 0.0376886 | MAP1LC3B;ACSL1;LPCAT3;SLC40A1;GCLM |
| Cellular senescence | 13/160 | 0.0377389 | MAP2K3;TGFB1;GADD45B;GADD45A;PIK3CB;MAPK14;HLA-E;GADD45G;CCNA1;MAPKAPK2;MCU;MAP2K6;MAPK3 |
| ECM-receptor interaction | 8/82 | 0.0377987 | GP9;ITGB4;ITGA2B;ITGA7;SDC1;GP1BA;LAMC1;THBS3 |
| Prostate cancer | 9/97 | 0.0378283 | NFKBIA;CASP9;IL1R2;TGFA;GRB2;PIK3CB;IKBKG;MMP9;MAPK3 |
| Autophagy | 11/128 | 0.0378376 | SH3GLB1;VMP1;MTMR3;DAPK2;PRKCD;ULK1;PIK3CB;CFLAR;CTSD;RAB7A;MAPK3 |
| Epithelial cell signaling in Helicobacter pylori infection | 7/68 | 0.03928 | LYN;NFKBIA;ADAM17;JUN;IKBKG;MAPK14;ATP6V1C1 |
| Phenylalanine metabolism | 3/17 | 0.0425463 | MAOB;MAOA;HPD |
| *Inhibited* | *Pathways* |  |  |
| Herpes simplex virus 1 infection | 262/492 | 1.37E-33 | ZNF175;ZNF607;ZNF606;ZNF727;AKT2;ZNF605;AKT3;ZNF846;AKT1;ZNF721;ZNF600;ZNF841;ZNF169;ZNF17;ZNF285;ZNF19;ZNF284;ZNF283;ZFP1;ZNF10;ZNF12;HCFC2;ZNF718;ZNF717;ZNF836;TP53;ZNF398;ZNF275;ZNF154;PIK3R3;PIK3R1;ZNF25;ZNF26;ZNF829;ZNF708;ZNF707;ZNF823;ZNF268;HLA-DQA2;HLA-DQA1;ZNF264;HLA-DRB5;ZNF141;ZNF383;ZNF382;ZNF140;ZNF34;EIF2AK4;ZNF37A;ZNF816;ZNF814;ZNF30;ZNF813;RBAK;ZNF136;ZNF257;ZNF256;ZNF135;HLA-DRB1;ZNF133;ZNF253;ZNF132;ZNF251;ZNF250;ZNF490;ZNF43;ZNF44;ZNF45;IKBKB;ZNF41;ZNF248;IKBKE;ZNF124;HLA-DPA1;JAK1;ZIK1;ZNF484;ZNF480;TMEM173;ZNF57;ZNF780B;ZNF780A;ZNF599;ZNF235;ZNF234;ZNF597;ZNF596;ZNF595;ZNF473;ZNF470;ZNF2;ZNF3;ZNF8;ZNF7;HLA-DMA;ZFP14;HLA-DMB;ZNF227;ZNF226;ZNF468;ZNF347;ZNF589;ZNF225;ZNF585B;ZNF224;ZNF587;ZNF343;ZNF101;ZNF585A;ZNF584;ZNF100;ZNF583;CD74;ZNF461;ZNF582;CARD9;ZNF79;ZNF33B;PPP1CA;ZNF74;HLA-DPB1;ZNF337;ZNF699;ZNF577;ZNF212;ZNF354C;ZNF211;ZFP28;ZNF354B;ZNF331;ZNF571;ZNF570;TNF;PPP1CC;ZNF81;ZFP30;ZNF84;ZNF208;ZNF85;ZNF569;ZNF689;ZNF568;ZNF567;ZNF566;HLA-DOA;ZNF565;ZFP37;MAP3K7;ZNF563;ZNF562;ZNF320;ZNF440;ZNF561;ZNF682;ZNF680;TSC1;ZNF90;ZNF91;ZNF439;ZNF317;ZNF559;SRSF2;SRSF3;ZNF558;ZNF557;ZNF436;SRSF5;SRSF6;ZNF555;SRSF7;ZNF675;ZNF554;SRSF8;ZNF432;ZNF431;ZNF551;ZNF793;ZNF430;ZNF792;ZNF671;ZNF550;ZNF670;ZNF791;ZNF790;ZNF549;ZNF669;ZNF548;ZNF547;ZNF426;ZNF546;ZNF786;ZNF302;ZNF544;ZNF543;ZNF785;ZNF300;ZNF420;ZNF783;EIF2B5;ZNF540;ZNF782;EIF2B4;EIF2B2;POU2F2;ZFP69;ZNF419;ZNF418;ZNF417;ZNF416;ZNF658;BCL2;ZNF778;HLA-DRA;TAB1;ZNF773;ZNF530;ZNF772;ZNF891;TRADD;FASLG;ZFP82;ZNF529;ZNF528;ZNF649;ZNF527;ZNF404;ZNF765;EIF2B1;ZNF764;ZNF761;TRAF2;ZFP90;ZNF519;IRF3;IFNG;TRAF3;ZNF517;TRAF5;ZNF879;ZNF514;ZNF510;BIRC3;ZNF195;SRC;SRSF1;NXF1;ZNF749;ZNF506;ZNF624;ZNF623;ZNF621;ZNF286A;ZNF184;ZNF181;ZNF180;MTOR;ZNF619;ZNF615;ZNF736;ZNF614;ZNF611;ZNF850 |
| Ribosome | 94/153 | 2.62E-18 | RPL4;MRPS17;RPL5;RPL30;RPL3;MRPS16;RPL32;RPL31;MRPS14;RPL34;MRPS11;RPLP1;MRPS12;RPLP0;RPL8;RPL10A;MRPL34;RPL9;MRPL35;MRPL32;RPL7;RPS15;MRPL4;RPS4X;RPS14;RPL7A;RPS17;RPS16;RPS19;RPL18A;RPS18;RPL36;RPL35;RPLP2;RPL38;RPL37;RPS11;RPL39;RPS10;RPS13;RPS12;RPL21;RPS7;RPS8;RPL23;RPS5;RPL22;RPS6;RPL13A;MRPS2;MRPS21;RPSA;RPS3A;MRPS7;MRPS6;MRPS5;MRPS9;RPL37A;RPL24;RPL27;RPL26;RPL29;UBA52;RPL12;MRPL19;RPL11;RPL36A;RPS27L;MRPL14;RPS15A;RPL14;RPS3;RPL13;RPL15;RPS2;RPL18;RPS27A;RPL17;RPL19;RPL41;MRPL27;RPL35A;RPL23A;MRPL21;MRPL30;RPS25;RPS28;RPS27;RPS29;RPL27A;RPS20;RPS21;RPS24;RPS23 |
| RNA transport | 80/165 | 1.09E-08 | CYFIP2;EIF4A2;POP5;EIF4A1;NUP107;NUP188;POP1;GEMIN2;RPP30;PHAX;NXT2;SUMO2;XPO5;EIF2B1;NDC1;NUP210;NCBP1;NUP133;NCBP2;ELAC2;PABPC4;ELAC1;THOC1;THOC3;THOC2;TRNT1;RANGAP1;SRRM1;THOC6;NUP93;EEF1A1;DDX39B;CLNS1A;RPP21;GEMIN4;GEMIN5;GEMIN6;NUP54;PABPC1;RPP25;GEMIN8;EIF4E2;NUP205;POM121;SEH1L;RBM8A;DDX20;NMD3;AAAS;NUP160;NXF1;NUP85;TGS1;EIF4EBP2;NUP88;NUP43;RPP14;RAE1;EIF4B;PAIP1;EIF2B5;EIF2B4;UBE2I;EIF2B2;NUP155;UPF3B;UPF3A;SNUPN;XPOT;EIF3G;NUP35;EIF3H;ACIN1;RNPS1;EIF3E;NUPL2;EIF3D;EIF4G2;EIF3A;EIF3B |
| Spliceosome | 63/134 | 1.46E-06 | TCERG1;RBM25;DDX46;PRPF19;USP39;PQBP1;EFTUD2;SNRPD2;SNRNP70;DHX15;DHX16;SNRPD3;CTNNBL1;AQR;NCBP1;NCBP2;THOC1;PRPF40B;THOC3;THOC2;PLRG1;WBP11;CDC40;PRPF4;RBMXL1;DDX39B;PRPF3;SRSF2;SRSF3;PPIE;PPIH;RP9;SRSF5;SRSF6;SNRNP200;SRSF7;SNRPA;SRSF8;DDX5;SF3B3;DHX8;RBM8A;SRSF1;PRPF8;PUF60;U2AF2;TRA2B;TRA2A;SF3B1;SRSF10;SF3A3;HSPA8;PRPF38A;CDC5L;LSM4;U2SURP;LSM8;LSM7;SNRNP40;LSM6;ACIN1;HNRNPA1L2;RBMX |
| RNA degradation | 39/79 | 3.57E-05 | ZCCHC7;DDX6;DIS3L;WDR61;TENT4A;ENO2;TOB2;TENT4B;ENO3;EDC3;EDC4;EXOSC6;EXOSC5;EXOSC10;EXOSC9;EXOSC8;DHX36;MTREX;EXOSC2;EXOSC1;HSPA9;TTC37;CNOT6L;DIS3;PABPC4;CNOT10;LSM4;PAN2;LSM8;CNOT6;PFKL;LSM7;CNOT7;LSM6;CNOT1;PABPC1;CNOT9;DCP1B;MPHOSPH6 |
| T cell receptor signaling pathway | 44/101 | 4.53E-04 | ITK;PIK3R3;CBLB;CD3G;PIK3R1;TNF;RASGRP1;CD3D;MALT1;IKBKB;MAPK9;PPP3CB;PPP3CC;AKT2;AKT3;AKT1;CTLA4;PAK6;FYN;PLCG1;ICOS;MAP3K7;HRAS;PAK4;PDPK1;NFATC3;NFATC2;NFATC1;VAV2;ZAP70;MAPK11;DLG1;CD4;CD40LG;IFNG;CDK4;LCK;CD28;PRKCQ;PDCD1;CD247;MAP3K14;CARD11;LAT |
| Th1 and Th2 cell differentiation | 40/92 | 8.44E-04 | CD3G;GATA3;CD3D;DLL1;IKBKB;MAPK9;PPP3CB;MAPK8;HLA-DMA;PPP3CC;HLA-DMB;TBX21;STAT4;PLCG1;HLA-DOA;HLA-DQA2;HLA-DQA1;HLA-DPA1;JAK1;IL12RB2;JAG2;HLA-DRB5;NFATC3;NFATC2;NFATC1;RUNX3;ZAP70;MAPK11;CD4;MAF;IFNG;LCK;IL2RA;IL2RB;HLA-DPB1;HLA-DRA;PRKCQ;CD247;LAT;HLA-DRB1 |
| Ribosome biogenesis in eukaryotes | 43/101 | 9.31E-04 | POP5;RBM28;POP1;NVL;RPP30;WDR3;HEATR1;NAT10;NMD3;SPATA5;NXT2;WDR43;NOL6;NXF1;EMG1;NOB1;TCOF1;BMS1;DROSHA;RIOK2;RIOK1;UTP14A;MDN1;UTP15;NOP56;WDR36;NOP58;UTP6;CSNK2A1;UTP4;IMP3;EFL1;CSNK2A2;WDR75;IMP4;GNL2;UTP18;GTPBP4;LSG1;GNL3L;SBDS;MPHOSPH10;RPP25 |
| Fanconi anemia pathway | 26/54 | 0.0011091 | MUS81;WDR48;REV3L;POLI;PMS2;POLK;ATRIP;POLH;FAAP100;TOP3B;CENPX;POLN;REV1;RPA1;FANCC;MLH1;FANCE;PALB2;FANCG;FANCF;RAD51C;ERCC4;ERCC1;FAN1;TELO2;ATR |
| Th17 cell differentiation | 44/107 | 0.0018969 | HSP90AB1;RORC;RORA;CD3G;GATA3;CD3D;IL27RA;IKBKB;MAPK9;PPP3CB;MAPK8;HLA-DMA;PPP3CC;HLA-DMB;TBX21;IL21R;PLCG1;HLA-DOA;HLA-DQA2;HLA-DQA1;HLA-DPA1;JAK1;SMAD4;HLA-DRB5;SMAD3;NFATC3;NFATC2;NFATC1;FOXP3;MTOR;ZAP70;MAPK11;CD4;IFNG;IL23A;LCK;IL2RA;IL2RB;HLA-DPB1;HLA-DRA;PRKCQ;CD247;HLA-DRB1;LAT |
| Nucleotide excision repair | 22/47 | 0.0040438 | LIG1;RFC1;CCNH;XPA;RPA1;RAD23A;GTF2H3;DDB2;CUL4A;DDB1;POLE4;ERCC3;ERCC4;ERCC1;POLD2;POLE3;ERCC2;ERCC8;ERCC5;ERCC6;MNAT1;POLE |
| RNA polymerase | 16/31 | 0.0041293 | ZNRD1;POLR3A;POLR2B;POLR3C;POLR1A;POLR3D;POLR1B;TWISTNB;POLR2D;POLR3E;POLR1C;POLR3F;POLR1E;POLR2H;POLR3K;POLR2K |
| mRNA surveillance pathway | 37/91 | 0.0052027 | DAZAP1;SMG1;RBM8A;RNMT;CSTF2;CSTF2T;MSI2;NXT2;SMG6;PPP1CC;NXF1;PPP2R1B;PPP2R5E;PCF11;CSTF1;SYMPK;CPSF6;RNGTT;NCBP1;CPSF1;NCBP2;PABPC4;CPSF3;UPF3B;PPP2R5D;UPF3A;PPP2R5C;SRRM1;WDR33;PPP1CA;NUDT21;DDX39B;WDR82;PPP2R2B;ACIN1;RNPS1;PABPC1 |
| Other types of O-glycan biosynthesis | 12/22 | 0.0071819 | LFNG;ST6GAL1;COLGALT2;POGLUT1;POFUT1;MFNG;POMT1;EOGT;GXYLT1;B3GLCT;PLOD3;OGT |
| Allograft rejection | 18/38 | 0.0076817 | CD86;CD40;HLA-DRB5;PRF1;FASLG;TNF;HLA-DMA;HLA-DMB;CD40LG;IFNG;CD28;HLA-DPB1;HLA-DRA;HLA-DOA;HLA-DQA2;HLA-DQA1;HLA-DRB1;HLA-DPA1 |
| Graft-versus-host disease | 19/41 | 0.0083365 | CD86;HLA-DRB5;PRF1;FASLG;KIR2DL1;TNF;HLA-DMA;HLA-DMB;IFNG;CD28;HLA-DPB1;HLA-DRA;KLRD1;KLRC1;HLA-DOA;HLA-DQA2;HLA-DQA1;HLA-DRB1;HLA-DPA1 |
| Mannose type O-glycan biosynthesis | 12/23 | 0.0112714 | B3GALNT2;FKRP;B4GAT1;POMT1;B3GAT1;CHST10;POMK;RXYLT1;POMGNT2;POMGNT1;LARGE2;FKTN |
| Asthma | 15/31 | 0.0114481 | CD40;HLA-DRB5;TNF;HLA-DMA;HLA-DMB;CD40LG;HLA-DPB1;FCER1A;HLA-DRA;HLA-DOA;MS4A2;HLA-DQA2;HLA-DQA1;HLA-DRB1;HLA-DPA1 |
| D-Glutamine and D-glutamate metabolism | 4/5 | 0.0230065 | GLUD1;GLS2;DGLUCY;GLS |
| Human T-cell leukemia virus 1 infection | 74/219 | 0.0280696 | ATF2;CD40;CRTC3;TRRAP;CRTC1;CD3G;ITGAL;ELK1;CD3D;TNF;ETS1;ELK4;IKBKB;PPP3CB;CDC23;PPP3CC;CREB3L4;CHEK2;MYC;AKT2;AKT3;AKT1;HLA-DOA;PRKACB;HRAS;HLA-DPA1;JAK1;TBP;KAT2B;KAT2A;ADCY9;CREB1;LCK;ANAPC4;VDAC1;ANAPC5;TP53;ANAPC1;ANAPC2;VAC14;ATF6B;PIK3R3;GPS2;PIK3R1;ADCY7;ANAPC10;MAPK9;HLA-DMA;MAPK8;HLA-DMB;BUB3;HLA-DQA2;HLA-DQA1;EGR2;HLA-DRB5;SMAD4;SMAD3;NFYB;TGFB3;NFATC3;NFATC2;NFATC1;DLG1;CD4;CDK4;IL2RA;CDC16;IL2RB;HLA-DPB1;HLA-DRA;ATM;MAP3K14;HLA-DRB1;ATR |
| Ubiquitin mediated proteolysis | 48/137 | 0.036571 | DET1;CUL7;UBE2D4;UBE3C;CUL5;CUL2;UBE3A;CBLB;UBE2Z;PRPF19;ANAPC10;RHOBTB2;CDC23;HERC2;HERC1;UBE2Q2;FBXO4;VHL;BTRC;SKP2;PIAS3;UBE2I;PPIL2;SMURF2;FBXW11;UBE2E3;HUWE1;UBE2E2;WWP1;UBE2G2;DDB2;DDB1;CUL4A;KLHL9;ITCH;CDC16;UBE2N;ANAPC4;UBA2;ERCC8;ANAPC5;BIRC6;TRIM37;STUB1;TRIM32;ANAPC1;ANAPC2;BIRC3 |
| Inflammatory bowel disease (IBD) | 25/65 | 0.039257 | RORC;RORA;GATA3;TNF;HLA-DMA;HLA-DMB;TBX21;IL21R;STAT4;HLA-DOA;HLA-DQA2;HLA-DQA1;HLA-DPA1;IL12RB2;HLA-DRB5;SMAD3;TGFB3;NFATC1;FOXP3;MAF;IFNG;IL23A;HLA-DPB1;HLA-DRA;HLA-DRB1 |
| Riboflavin metabolism | 5/8 | 0.0420706 | FLAD1;RFK;ACP5;ENPP3;ACP1 |
| Base excision repair | 14/33 | 0.0493072 | LIG1;OGG1;APEX2;LIG3;UNG;POLE4;NEIL2;NTHL1;TDG;POLD2;POLE3;POLL;POLE;MUTYH |
| Intestinal immune network for IgA production | 19/48 | 0.0504455 | CD86;CD40;HLA-DRB5;ITGA4;TNFSF13;HLA-DMA;HLA-DMB;CD40LG;CD28;HLA-DPB1;HLA-DRA;ICOS;HLA-DOA;HLA-DQA2;MAP3K14;CCL28;HLA-DQA1;HLA-DRB1;HLA-DPA1 |

**Proportion = Differentially Expressed Genes/ Total No of Genes in Pathwa***y**

Supplemental Table S3. Univariate validation of the 18 selected genes contributing to the Weighted Gene Expression Risk Score (WGERS).

|  | **Training** | | | | | | | **Validation** | | | |
| --- | --- | --- | --- | --- | --- | --- | --- | --- | --- | --- | --- |
| **Gene** | **Risk score**  **coefficient** | **mean** | **SD** | **AUC** | **Log Fold change** | **p-value** | **BH FDR** | **AUC** | **Log Fold change** | **p-value** | **BH FDR** |
| *Intercept* | *-1.013* |  |  |  |  |  |  |  |  |  |  |
| AATBC | -0.213 | 3.781 | 1.261 | 0.918 | -1.624 | 4.30E-07 | 2.96E-05 | 0.710 | -0.584 | 2.99E-04 | 3.37E-04 |
| AC104809.2 | -0.187 | 1.434 | 1.554 | 0.967 | -1.598 | 1.67E-08 | 2.96E-05 | 0.690 | -0.050 | 2.62E-04 | 3.27E-04 |
| CD4 | -0.211 | 7.459 | 1.183 | 0.964 | -1.689 | 2.07E-08 | 2.96E-05 | 0.874 | -1.859 | 1.23E-10 | 2.22E-09 |
| CDKN1C | -0.191 | 2.634 | 1.451 | 0.924 | -1.892 | 2.92E-07 | 2.96E-05 | 0.711 | -0.661 | 2.73E-04 | 3.27E-04 |
| CLEC10A | -0.205 | 3.267 | 1.203 | 0.909 | -1.719 | 7.61E-07 | 3.17E-05 | 0.842 | -1.774 | 3.69E-09 | 2.49E-08 |
| CSF1R | -0.213 | 7.195 | 1.269 | 0.914 | -1.757 | 5.73E-07 | 2.96E-05 | 0.834 | -1.414 | 9.14E-09 | 3.29E-08 |
| IL18R1 | 0.195 | 6.157 | 1.497 | 0.914 | 2.061 | 5.73E-07 | 2.96E-05 | 0.770 | 1.883 | 3.43E-06 | 5.50E-06 |
| MCEMP1 | 0.191 | 6.200 | 1.377 | 0.909 | 1.873 | 7.61E-07 | 3.17E-05 | 0.804 | 1.373 | 1.58E-07 | 3.16E-07 |
| MS4A14 | -0.205 | 3.226 | 1.158 | 0.921 | -1.685 | 3.54E-07 | 2.96E-05 | 0.721 | -0.679 | 1.38E-04 | 1.91E-04 |
| NEURL1 | -0.217 | 2.133 | 1.149 | 0.958 | -1.806 | 3.16E-08 | 2.96E-05 | 0.838 | -0.471 | 5.54E-09 | 2.49E-08 |
| NR4A1 | -0.199 | 2.477 | 1.175 | 0.927 | -1.613 | 2.40E-07 | 2.96E-05 | 0.769 | -0.672 | 3.67E-06 | 5.50E-06 |
| PID1 | -0.206 | 2.502 | 1.473 | 0.905 | -2.481 | 1.01E-06 | 3.35E-05 | 0.827 | -1.432 | 1.74E-08 | 5.22E-08 |
| SLC51A | 0.188 | 1.422 | 1.457 | 0.918 | 1.627 | 4.30E-07 | 2.96E-05 | 0.793 | 0.987 | 4.58E-07 | 8.25E-07 |
| SOCS3 | 0.198 | 7.045 | 1.110 | 0.926 | 1.679 | 2.65E-07 | 2.96E-05 | 0.656 | 0.529 | 7.25E-03 | 7.25E-03 |
| TGFBI | -0.202 | 6.640 | 1.490 | 0.924 | -1.874 | 2.92E-07 | 2.96E-05 | 0.840 | -1.579 | 4.73E-09 | 2.49E-08 |
| TPPP3 | -0.209 | 2.120 | 1.239 | 0.942 | -1.878 | 8.85E-08 | 2.96E-05 | 0.817 | -1.395 | 4.95E-08 | 1.11E-07 |
| ZDHHC19 | 0.193 | 3.175 | 2.206 | 0.906 | 2.780 | 9.18E-07 | 3.28E-05 | 0.692 | 1.046 | 9.25E-04 | 9.79E-04 |
| ZNF703 | -0.220 | 2.700 | 1.337 | 0.935 | -2.227 | 1.46E-07 | 2.96E-05 | 0.820 | -0.636 | 3.68E-08 | 9.47E-08 |

The first two genes were missing in the Validation Data, and were imputed via multiple linear regression based on the other 16 CPM-normalized log gene expression values, with adjusted R^2^ = 95.6% (AATBC) and 90.2% (AC104809.2) in the Training Data.

All 18 genes were differentially expressed (FDR < 0.01, correcting for the 18 comparisons) in the same direction but with uniformly lower univariate AUC in the Validation Data.

The risk score coefficients were designed to apply to Z-score standardized CPM-normalized gene expression values, with mean and SD estimated from the Training Data.

Fold changes were computed as the Hodges-Lehmann shift in log_2_(1 + CPM normalized gene expression), a robust location measure consistent with Wilcoxon.

p-values were based on the Wilcoxon test (which targets AUC), with Benjamini-Hochberg (BH) False Discovery Rates (FDR) corrected for all 14,228 genes in the Training Data but only the selected 18 genes in the Validation Data
